# Serum α-Klotho levels correlate with episodic memory changes related to cardiovascular exercise in older adults

**DOI:** 10.1101/2020.01.16.908913

**Authors:** Andreas Becke, Anne Maass, Michael R. Kreutz, Emrah Duezel

## Abstract

Aerobic exercise is a potential life-style intervention to delay cognitive decline and neurodegeneration. Elevated serum levels of the anti-aging protein α-Klotho (αKL) are a potential mediating factor of exercise benefits on cognition. Here, we examined in older adults how exercise-related changes of αKL levels in serum relate to changes in cerebral blood flow (CBF), hippocampal volumes and episodic memory. We analyzed data from a previously published intervention study in which forty cognitively healthy subjects were pseudo-randomly assigned to either a cardiovascular exercise group (treadmill training, n=21) or control group (indoor progressive-muscle relaxation/stretching, n=19). 3-Tesla gadolinium perfusion imaging was used to track hippocampal CBF changes and high resolution 7-Tesla-T1-images were used to track hippocampal volume changes. Changes in episodic memory performance were measured using the complex figure test (CFT). Longitudinal changes were compared between groups and analyzed with a multiple linear regression approach. CFT and hippocampal volume changes significantly predicted changes in serum αKL levels. For CFT, this effect was found in the exercise but not the control group. Collectively the data suggest that αKL level increases induced by exercise can be associated with improved hippocampal function in older adults.

## Introduction

Physical exercise is an important moderator preserving and improving cognitive function in the aging human brain (Colcombe et al., 2003; Kelly et al., 2014). The *klotho* gene was originally identified in mice as a putative age-suppressing gene that extends life span when overexpressed (Kuro-o et al., 1997; Kurosu et al., 2005). It induces complex phenotypes resembling human premature aging syndromes when disrupted (Kuro-o et al., 1997; Kurosu et al., 2005). Klotho is most prominently expressed in the kidney. Variations in the *klotho* gene are associated with both life extension and increased cognition in human populations. The Klotho-VS polymorphism in humans for instance promotes cognition by increasing serum levels of secreted αKL (Cararo-Lopes et al., 2017). The absence of Klotho in mice causes cognitive impairment, whereas increasing Klotho improves hippocampal-dependent memory (Dubal et al., 2014, 2015). Since elevating Klotho serum levels in mice enhances cognition (Dubal et al., 2014, 2015; Leon et al., 2017), strategies that increase Klotho activity or simulate its function may therefore improve cognition at different life stages and, possibly, even under pathological circumstances. Recent evidence suggests that Klotho is a regulator of postnatal neurogenesis, affecting neural stem cell proliferation and maturation sufficient to impact hippocampal-dependent spatial memory function (Laszczyk et al., 2017). Premature aging of the *klotho*-deficient hippocampal neurogenic niche is evident by a reduced number of neural stem cells, decreased proliferation, and impaired maturation of immature neurons (Laszczyk et al., 2017). Conversely, 6-months-old KL-overexpressing mice exhibit increased numbers of neural stem cells, increased proliferation, and more immature neurons with enhanced dendritic arborization. In addition, studies in humans indicate anti-aging features of αKL, for example, by cardiovascular protection (Marcais et al., 2017; Matsubara et al., 2014; Semba et al., 2011).

However, along with higher age lower levels of cerebral Klotho levels were found in rhesus monkeys (Duce et al., 2008; King et al., 2012) and humans (Yamazaki et al., 2010). With regard to potential therapeutic features, it has been therefore of great interest to understand how Klotho levels can be increased in humans. Physical exercise has been already shown to moderate or induce the release of neurotrophic and vascular growth factors, like BDNF, IGF-1 and VEGF (e.g. Maass et al., 2016; for review see also Cotman et al., 2007; Voss et al., 2013) as well as proteins mediating their functionality, like Cathepsin-B (Moon et al., 2016). Indeed a 12-week regular moderate aerobic exercise training in older women can increase the level of αKL in serum samples (Matsubara et al. 2014) and αKL has also been shown to be elevated in serum directly after a single bout of 20 min exercise at maximum intensity (Santos-Dias et al., 2017). Currently, it is still unclear whether beneficial effects of exercise on hippocampal memory functions are related to increased levels of peripheral αKL. We therefore tested the hypothesis that exercise-induced benefits in cognitive functions in older adults are related to αKL-induced structural and perfusion changes in the hippocampus.

## Methods

### Participants and experimental design

In a previously published (Maass et al., 2015), controlled 3-month intervention trial, 40 sedentary healthy older adults (mean age = 68.4 ± 4.3 years, 55% females) were randomly assigned to either aerobic exercise training group (N=21) or control (relaxation) group (N=19), matched by age, gender, cognitive baseline measures (verbal learning score) and body mass index. Study details can be found in Maass et al. (2015). All subjects received monetary compensation and signed written informed content of the study, which was proved by the ethics committee at the Otto-von-Guericke University Magdeburg.

Here we used remaining blood samples from this study to assess the relationship between exercise induced neurovascular plasticity in the brain and serum levels of αKL.

### Exercise intervention and spiroergometric testing

The cardiovascular exercise intervention regime consisted of treadmill training 3x per week for 30-40 minutes at target heart rate, starting at 65% of heart rate maximum and increasing by 5% in steps for 4 weeks. The intensity of the training was monitored via chest heart rate monitors controlling automatic speed and incline of the treadmills. Subjects in the control group perceived in the same amount of time per week a relaxation technique training (Jacobson, 1929) to match social interactions, schedule, and motivation between groups (for details see Maass et al., 2015).

Consumption of oxygen at ventilatory threshold (VO2-VT) was assessed by graded maximal exercise testing on recumbent cycle ergometer. From an initial resistance of 20 Watt the intensity increased every 2 min by 20 Watt. Subjects were asked to keep a constant cadence at 60-70 repetitions/min. This occurred until fulfilling exhaustion criteria were met, e.g. a respiratory exchange ratio (RER) higher than 1.1, heart rate reached the calculated individual maximum or cardiovascular abortion criteria appeared via pulmonary cardiac online observation.

At the end of each training session subjects in the training group were asked to rate their perceived exertion (RPE_BORG_) on the CR-10 scale (Borg, 1998) in steps of one from 6 (no exhaustion at all) to 20 (maximum exhaustion). Average values over all training sessions were calculated to indicate individual intensity levels.

To measure other physical activity next to the exercise intervention regime given by the study, subjects were asked to answer the International Physical Activity Questionnaire (IPAQ) by the end of each week. Metabolic rates (kcal/kg/h) were calculated in sum over the duration of the intervention in respect to the intensities of the physical activities (metabolic index) in accordance to the IPAQ scoring protocol (see [www.ipaq.ki.se] revised version, 2005).

### Blood biochemistry

Fasting blood samples were collected from antecubital vein at baseline before the cardiovascular examination and post intervention within 3 days of MRI scans. The samples were cooled at 4°C before centrifuged. Serum samples were aliquoted and stored at −20°C. The aliquots were used to analyze blood for PDGF, IGF/IGF-Bps, cortisol (Maass et al., 2016) and soluble αKL by ELISA (IBL international, sensitivity 6.15 pg/ml, range 93.75 – 6000 pg/ml).

### Cognitive measures

An extensive neurological test battery was conducted before and after the intervention of the study (for full details see Maass et al., 2015), including the Complex Figure Test (CFT); containing elements for pattern separation memory performance tested in early recall, late recall and recognition (Strauss et al., 2006). For the baseline measures the Rey-Osterrieth Complex Figure (ROCF) was administered, whereas the Modified Taylor Complex Figure (MTCF) served as the post-intervention measure. MTCF is a valid parallel version of the ROCF avoiding implicit learning for repeated testing sessions (Casarotti et al., 2014). The compound method of MTCF and ROCF is further referred as complex figure test (CFT).

CFT consists of different geometric patterns, which can be divided into 18 elements. The test procedure is as follows: First, the figure has to be copied as detailed as possible (copy trial). After a 3-min (early recall) and a 30-min delay interval (late recall), participants are asked to reproduce the figure as accurate as possible from memory. Recognition of the CFT is tested immediately after the late recall trial. The recognition trial presents 12 of the 18 scoring elements of the before shown complex figure, along with 12 new elements (from the other version) that serve as lures and participants have to indicate which items they recognize from the original figure. The recognition trial requires discrimination between highly similar objects, and thereby poses high demands on hippocampal pattern separation.

### Hippocampal subfield volume measures

To measure changes in hippocampus volumes, segmentation of hippocampal regions was performed manually on the 7T (Siemens, Erlangen, Germany) high-resolution MPRAGE images (0.6 mm isotropic, TE = 2.8 ms, TR = 2500 ms, TI = 1050 ms, flip angle = 5°, acquisition time 14:00 min). The left and right hippocampus was further subdivided into head, body, and tail sections (for further details on the segmentation protocole see Maass et al., 2015).

### Brain Perfusion measures

Gadolinium contrast-based perfusion imaging at 3T (Siemens Magnetom, Verio, 32-channel head coil) was used to measure changes cerebral blood volume (CBV) in resting condition. High-resolution (partial) perfusion-weighted images were acquired with slice alignment parallel to the hippocampal main axis (TR/TE = 1500/30 ms, 1.6 mm in-plane resolution, 3 mm slice thickness, 20 slices with 10% gap) and quantitative perfusion maps CBV were calculated. Additionally, general (non-hippocampal) gray-matter perfusion was calculated. The perfusion analyses are described in detail in Maass et al. (2015).

### Statistical analyses

Variables of interest were αKL in serum, oxygen consumption in a cardiovascular fitness test (VO2-VT), hippocampus volume (HC_VOL_), blood perfusion (blood volume and blood flow) for whole brain grey matter (GM_CBV/CBF_) and the hippocampus (HC_CBV/CBF_)) and recognition memory (CFT). For statistical evaluation in SPSS version 25.0 (IBM Corp.) was used. Repeated-measures ANOVA were calculated with intervention group as between-subject factor and time (pre vs. post) as within-subject factor.

To examine associations between measures, percentage changes were calculated from pre-to post measure data for all predictor variables and adjusted for age, gender and outside temperature change (to account for season effects) over study visits. Automatic linear regression analysis was used to test predictors (adjusted models) of αKL. First we tested a baseline model to identify the multivariable predictors of αKL. In a post-hoc automatic linear regression analysis, changes of hippocampus subfield volumes were tested separately as predictors of αKL. Next, we designed a multivariable model, including the covariates that had emerged as significant ones in the multivariable analysis and compared them between groups of intervention. Significant predictors were then followed up using in Pearson correlation coefficients. Statistical significance was set a priori at p < .05 for all comparisons.

## Results

Repeated-measure ANOVAs revealed a significant group (exercise versus control) by time (before and after exercise) interaction for VO2-VT (F(1,36) = 5.83; p = .022). Other outcome measures such as αKL, HC volume, CBV, CBF, and CFT did not show a group by time interaction (all p > .05). For an overview of mean percentage changes of all measures included in the analysis see figure 1.

**Figure 1.**
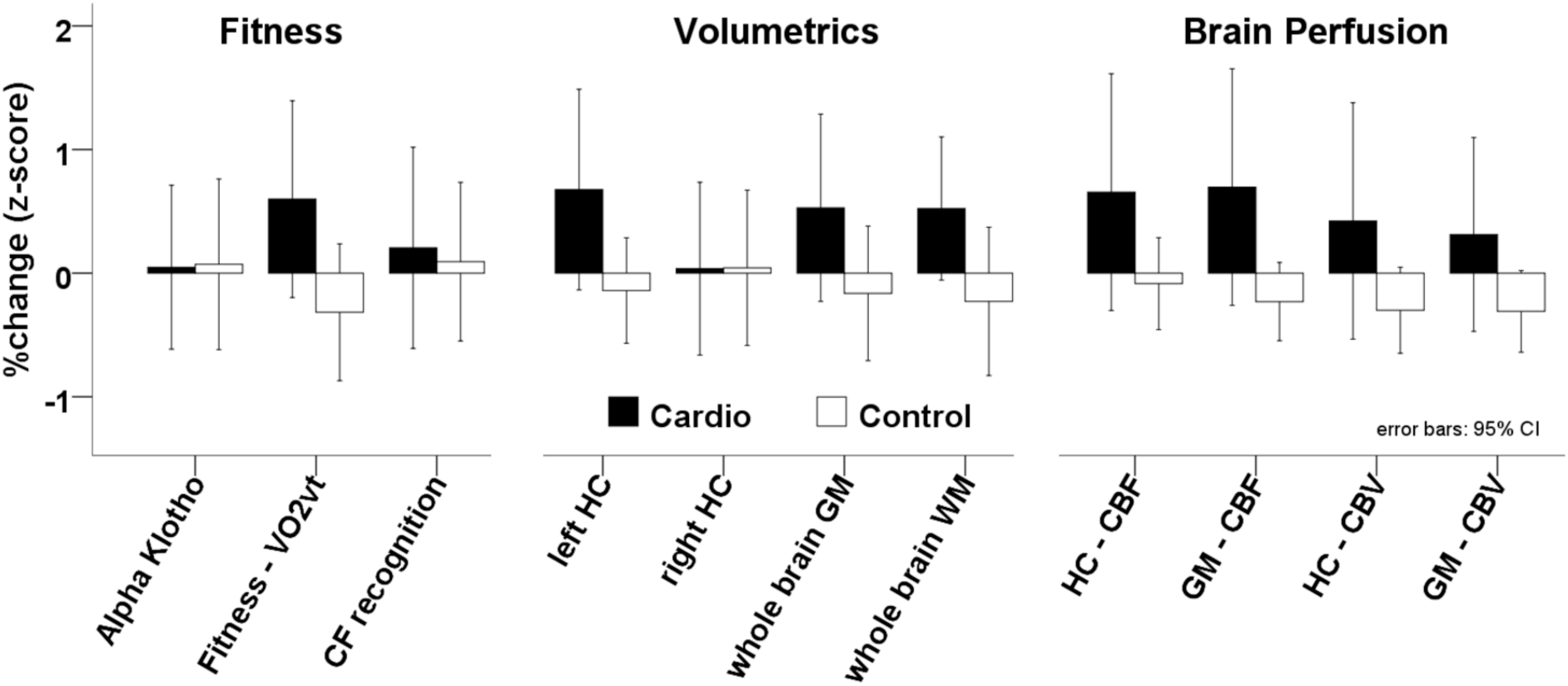
Normalized (z-scores [over all subjects])) % change values (post-minus pre-intervention) for αKL levels (Serum), oxygen volume at ventilatory threshold (VO2vt), complex figure recognition score (CF), left and right hippocampus (HC) volume, volumetric measures of grey matter without HC, white matter volume, cerebral blood flow (CBF) and cerebral blood volume (CBV). Filled bars = cardiovascular training group; open bars = control group.

An automatic linear regression model with changes αKL as independent variable (dependent variables: VO2-VT, total HC_VOL_, GM_CBV/CBF,_ HC_CBV/CBF_, CFT performance) revealed a model fit with R^2^_adj_ = .244 (corrected Akaike Information Criterion= 242.14). Significant predictors for αKL were CFT (t = 2.84; p = .008) and HC volume (t = 2.66; p = .012). HC_CBF_ also survived as a predictor for αKL but this was not significant (t = −1.23; p = .228).

To illustrate the results indicated by the linear regression analysis, table 1 provides an overview of correlations between αKL and outcome parameters of interest. For hippocampal volume, the table shows correlations separately for head, body and tail. As expected from the regression analysis, there are significant correlations between αKL and HC volumes and αKL and cognition (CFT). Separate assessments for each group show that the Cardio group drives both correlations.

**Table 1.** Pearson correlations of αKL pre-to-post changes (%) with CFT (complex figure recognition), bilateral hippocampus volume (HCbilateral), hippocampus section volumes (head and tail) and cerebral blood flow for bilateral hippocampus (HCCBF); displayed separately in rows for the whole study sample (All), the cardiovascular training group (Cardio) and control group (Control). *p<.05; **p<.01 (2-tailed)

|  |  | Cognition | Hippocampus Volumes |  |  | Hippocampus Perfusion |
| --- | --- | --- | --- | --- | --- | --- |
| | | CFT | $HC_{bilateral}$ | Head | Tail | $HC_{CBF}$ |
| **ALL** | R | .43** | .368* | .375* | .339 | .159 |
|  | p | .008 | .045 | .041 | .066 | .401 |
|  | N | 36 | 30 | 30 | 30 | 30 |
| **Cardio** | R | .563* | .416 | .497 | .371 | .179 |
|  | p | .012 | .139 | .071 | .191 | .506 |
|  | N | 19 | 14 | 14 | 14 | 16 |
| **Control** | R | .280 | .35 | .294 | .334 | -.115 |
|  | p | .277 | .184 | .269 | .206 | .684 |
|  | N | 17 | 16 | 16 | 16 | 15 |

Looking closer at the section volumes of the hippocampus, this effect is significant for the hippocampus head (HH_VOL_) only. The correlation of HH_VOL_ and αKL is marginally significant when considering the Cardio group only. Previous analyses of the dataset had revealed that the hippocampus perfusion changes are highly correlated to hippocampus volume changes (see Maass et al., 2015). However, αKL and hippocampal perfusion changes (HC_CBF_) do not show a significant correlation.

**Figure 2.**
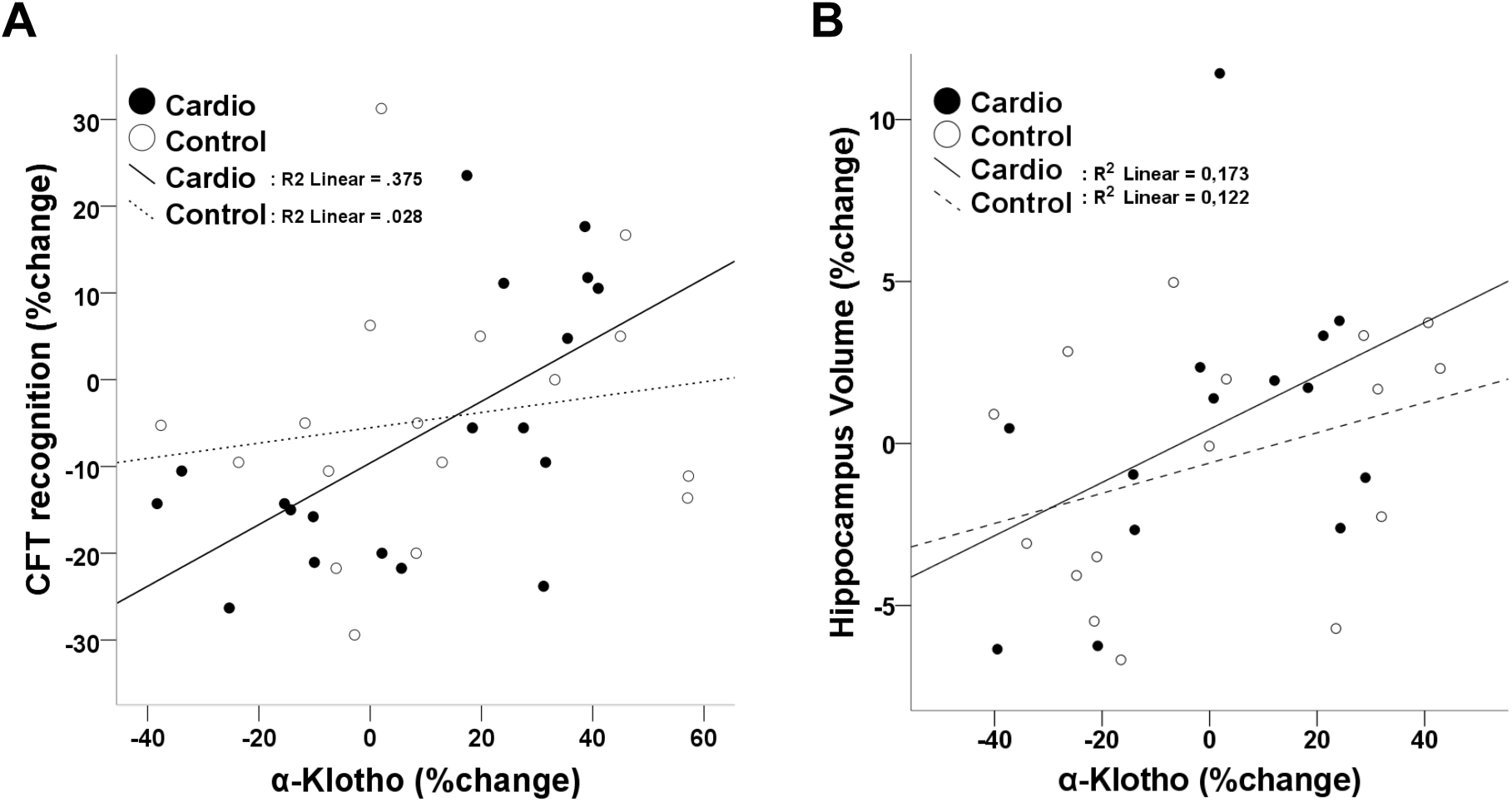
Scatterplots show correlations for changes in α-Klotho serum levels (after a 3 month cardiovascular exercise intervention in older adults) with percentage changes in memory recognition accuracy (A; delayed recall for Rey-complex-figures (CTF)) and changes in hippocampal volumes (B; bilateral volumes obtained from manually segmented 7Tesla-MPRAGE 0.6mm isotropic images).

The secondary outcome measure, mean RPE_BORG_ values, which were averaged over all training sessions, correlated positively with αKL percent changes in the training group (Pearson-R = .486, p = .030, N = 20) but no correlation with VO2_VAT_ (Pearson-R = -.135, p = .593). No rated perceived exertion was assessed for the control group after relaxation training sessions. Given that most commonly high training intensities related to high RPE_BORG_ values in training sessions with long durations or with high intensities, these data suggest that a more intense training relates to higher serum level increases in αKL.

## Discussion

The relationship between physical exercise related changes in αKL serum levels, changes in hippocampal volume and hippocampus-dependent memory have not yet been investigated in humans, to the best of our knowledge. Here we report that aerobic exercise was associated with a significant correlation of αKL changes in serum levels and a hippocampus-dependent memory task, whereas a non-exercise control intervention was not. Also, hippocampus volume changes were related to αKL changes in the aerobic exercise but not the control group.

These results are in agreement with former studies that demonstrated that exercise increases αKL levels (Matsubara et al., 2014, Santos-Dias et al., 2017) and cognitive improvement induced by αKL in animal studies (Dubal et al., 2014, 2015). As indicated by other animal studies (Leon et al., 2017), despite impermeability to the blood brain barrier, the administration of αKL protein fragment in the periphery can increase cognitive functions with neural resilience. These animal studies raise the possibility that the correlation of αKL changes and HC_VOL_ changes could be due to indirect effects of αKL on neuroplasticity. Collectively, the data support the possibility that αKL serum level increases can improve hippocampal plasticity and could thereby impact cognition and hippocampal function in aged persons. This effect of αKL on HC_VOL_ is in line with findings that Klotho is an important factor for the maturation of hippocampal neurons with impact on hippocampal-dependent spatial memory function (Laszczyk et al., 2017).

In a previous report of data from the same sample and study, we had reported that exercise induced hippocampus volume changes correlate to hippocampal perfusion changes (Maass et al., 2015). We found now that changes in αKL levels were not related to these changes in hippocampus perfusion. Nevertheless, it is worth pointing out that animal studies showed that blocking the genetic expression of Klotho can impact brain regulation for optimal blood pressure functionality. Furthermore, it was shown that silencing brain Klotho relates to the functional regulation of blood pressure via sympathetic nervous activity (Wang and Sun, 2010).

We also analyzed whether changes in αK levels were related to the subjectively experience exercise intensity. We indeed found a positive correlation of the mean perceived exertion after training and changes in αK levels during the course of the study. Although the training regime was conducted with respect to the individual fitness level, based on the heart rate at ventilatory threshold levels, the subjective responses indicate that the individual physical response to the training differs. Although a direct proof that αKL is being altered by the exercise intervention itself was is difficult to provide, regression analyses indicate that αKL related HC-Volume changes were dependent on respiratory fitness changes (VO2-VT).

These data indicate that the exercise intensity is crucial for the expression of αKL. The form of exercise might be crucial and a certain intensity may be required to induce exercise-dependent αKL release. Therefore, it is also plausible that the fitness marker of VO2-VT is not the strongest predictor for αKL release. As all subjects trained at their individual submaximal heart rate optimum and the variance in VO2-VT change was kept in a relatively small range. In line with our differential effects between groups, other studies already revealed the impact of exercise in human with Klotho levels being elevated in trained and healthy subjects compared to untrained subjects (Saghiv et al., 2016). A missing main effect of αKL between groups in our study is likely due to a lack of power as indicated by the high variance of αKL changes in the control group.

## Acknowledgments

We thank the German Center for the German Center for Neurodegenerative Diseases and the Otto-von-Guericke University (IKND) for funding this study and the Leibniz Institute for Neurobiology for providing access to their 7 T MR scanner.

